# Dense but not alpha granules of platelets are required for insulin secretion from pancreatic β cells

**DOI:** 10.1101/2024.09.23.614422

**Authors:** Katarzyna Kolczyńska-Matysiak, Till Karwen, Mona Loeffler, Izabela Hawro, Toufic Kassouf, David Stegner, Grzegorz Sumara

**Affiliations:** Nencki Institute of Experimental Biology, Polish Academy of Sciences, 3 Pasteur Street, 02-093 Warszawa, Poland; Rudolf-Virchow-Zentrum. Center for Integrative and Translational Bioimaging, University of Würzburg, 97080 Würzburg, Germany; Institute of Experimental Biomedicine I, University Hospital Würzburg, 97080 Würzburg, Germany

**Keywords:** platelet, insulin secretion, pancreatic β cell, platelet degranulation, dense granule secretion

## Abstract

**Objectives:** Platelets, originally described for their role in blood coagulation, are now also recognized as key players in modulating inflammation, tissue regeneration, angiogenesis, and carcinogenesis. Recent evidence suggests that platelets also influence insulin secretion from pancreatic β cells. The multifaceted functions of platelets are mediated by the factors stored in their alpha granules (AGs) and dense granules (DGs). AGs primarily contain proteins, while DGs are rich in small molecules, and both types of granules are released during blood coagulation. Specific components stored in AGs and DGs are implicated in various inflammatory, regenerative, and tumorigenic processes. However, the relative contributions of AGs and DGs to the regulation of pancreatic β cell function have not been previously explored.

**Methods:** In this study, we utilized mouse models deficient in AG content (neurobeachin-like 2 (*Nbeal2*) -deficient mice) and models with defective DG release (*Unc13d*-deficiency in bone marrow-derived cells) to investigate the impact of platelet granules on insulin secretion from pancreatic β cells.

**Results:** Our findings indicate that AG deficiency has little to no effect on pancreatic β cell function and glucose homeostasis. Conversely, mice with defective DG release exhibited glucose intolerance and reduced insulin secretion. Furthermore, *Unc13d*-deficiency in hematopoietic stem cells led to a reduction in adipose tissue gain in obese mice.

**Conclusions:** Obtained data suggest that DGs, but not AGs, mediate the influence of platelets on pancreatic β cells, thereby modulating glucose metabolism.

## 1. INTRODUCTION

Platelets are small anucleated cells originating from megakaryocytes, primarily known for their role in blood coagulation [1]. Upon vascular injury, platelets adhere to the exposed extracellular matrix proteins, sealing the injured surface, which leads to their activation and facilitates platelet degranulation [1]. Three major types of granules can be released from platelets: α-granules (AGs), dense granules (DGs), and lysosomes [2]. AGs and DGs are formed by distinct mechanisms and contain specific cargo. AGs are filled with around 300 proteins that play a role in platelet adhesion and also modulate inflammatory processes, wound healing, and angiogenesis [3]. In contrast, DGs contain small molecules, including serotonin, ATP, ADP, and various cations [2]. Generally, the release of DG content propagates and accelerates platelet aggregation through auto- and paracrine mechanisms [2].

In humans, loss of function mutations in *Neurobeachin-like 2* (*NBEAL2*) results in the formation of platelets depleted from AGs [3-5]. This condition, known as Grey platelet syndrome, is characterized by prolonged bleeding, mild to moderate thrombocytopenia, platelets lacking visible AG, and an almost complete loss of AG cargo [3, 4]. Mice deficient for *Nbeal2* replicate all the phenotypes observed in humans [3, 5]. Despite defects in neutrophil degranulation, patients and rodents lacking NBEAL2 or expressing its non-functional variants, primarily present platelet-specific abnormalities [6].

Degranulation of platelets requires the induction of the exocytotic machinery, with Munc13-4 (encoded by *UNC13D*) being an essential component of this pathway [7]. Released agonists further promote platelet activation, degranulation, and the functional upregulation of GPIIb/IIIa to bind fibrinogen and other multimeric ligands, resulting in platelet aggregation [8]. Platelets lacking Munc13-4 are unable to secrete their DG content [9].

Although platelets are primarily responsible for preventing excessive bleeding upon injury, recent decades have highlighted their central role in modulating inflammatory and regenerative processes [10]. Recently, a cross-talk between platelets and pancreatic β cells has been described. A fraction of platelets resides at the endothelium of pancreatic islets. Moreover, platelet activity is stimulated by glucose and pancreatic β-cell-derived factors. In turn, platelets secrete lipid-based substances (including 20-Hydroxyeicosatetraenoic acid – 20-HETE) that stimulate insulin secretion to maintain glucose homeostasis. Consequently, mice depleted of platelets or presenting defects in platelet adhesion or activation show lower circulating insulin levels and glucose intolerance [11]. However, it is not clear if defective AG or DG secretion in platelets contributes to the development of glucose intolerance.

In this study, we utilized mouse models of AG deficiency (*Nbeal2*^*-/-*^ mice) and animals with severe defects in DG degranulation (*Unc13d*^*BMC-/-*^ mice) to test the contribution of both types of platelet granules to the maintenance of glucose homeostasis. We showed that deficiency in platelet AGs does not affect insulin secretion or glucose tolerance. However, animals with defects in DG release present glucose intolerance associated with defective glucose-stimulated insulin secretion. Moreover, obese mice with *Unc13d*-deficiency in bone marrow-derived cells exhibit decreased glucose tolerance and reduced adiposity. In conclusion, the data that was obtained suggests that DGs, but not AGs, are required for platelet-mediated insulin secretion.

## 2. METHODS

### Animal models

All experiments involving mouse models were approved by the district government of Lower Franconia (Bezirksregierung Unterfranken, references 55.2.2-2532-2-1296, 55.2-2532-2-999, 55.2-2532-2-435, and 55.2-2532-2-746) and the local ethics committee of Warsaw (I Lokalna Komisja Etyczna ds. Doświadczeń na Zwierzętach, references 1054/2020 and 1074/2020). Animals were maintained under specific pathogen□free conditions in the animal facilities of the Rudolf Virchow Centre and the Nencki Institute of Experimental Biology on a 12:12 light-dark cycle with free access to food and water. *Nbeal2*-deficient mice were previously described [3, 5]. Similarly, *Unc13d*-deficient mice were generated as described previously [9].

### Bone marrow transplantation

C57BL/6JRj male mice (4-week-old) were irradiated with 10 Gy using a radiation device (Faxitron). The animals were reconstituted by intravenous injection with 4 × 10^6^ bone marrow cells from the femur and tibia of donor male mice (*Unc13d*^*−/−*^ or littermate *Unc13d*^*+/+*^). Following transplantation, mice were treated with 2 mg/ml neomycin-sulfate (Sigma-Aldrich) in drinking water.

### Metabolic tests

For glucose tolerance tests, fasted animals were injected intraperitoneally with 2 g of glucose per kg of body weight in saline solution, and glucose levels were measured using an automated glucometer (Accu-Chek, Roche). Similarly, an insulin tolerance test was performed on fasted animals injected with 0.5 U of insulin (Sigma) per kg of body weight. For glucose-stimulated insulin secretion tests, fasted mice were injected with 3 g of glucose per kg of body weight, and blood samples for insulin measurements were obtained from the tail vein. Insulin levels were measured by ELISA (Crystal Chem).

Oxygen consumption, CO2 dissipation, food intake, and voluntary movements were analyzed using the Phenomaster system from TSE systems as previously described [12]. Body composition was analyzed using a minispec body composition analyzer (Bruker). Histological analyses of the pancreas were performed as described before [11]. Total pancreatic insulin and glucagon content were measured as described before [13].

### Statistics

Results are presented as mean values ± standard error of the mean (SEM). TheMann-Whitney test was used for the analysis of the significance of the data. P values lower than 0.05 were considered statistically significant. Animals were assigned to the groups randomly.

## 3. RESULTS

### 3.1. Nbeal2-deficiency does not affect glucose homeostasis

To determine the role of AGs in the regulation of insulin secretion and glucose homeostasis, we utilized NBEAL2-deficient mice *(Nbeal2*^-/-^), which exhibit an almost complete lack of AGs in platelets [5]. Similar to other mouse models with defects in platelet content, activation, or adhesion [14], we performed all metabolic assays in young adult mice (8-12 weeks old) fed a standard chow diet. Ten-week-old *Nbeal2*^-/-^ mice presented unaltered body weight (Figure 1A). Likewise, these mice showed similar glucose tolerance compared to control animals (Figure 1B). However, *Nbeal2*^*-/-*^ mice exhibited slightly, but significantly reduced fasting glucose levels (Figure 1B). Nonetheless, glucose-stimulated insulin secretion and peripheral insulin sensitivity were not altered by the deletion of *Nbeal2* (Figure 1C and D). Altogether, these data indicate that AG deficiency in platelets only marginally affects glucose homeostasis and does not alter insulin secretion.

**Figure 1.**
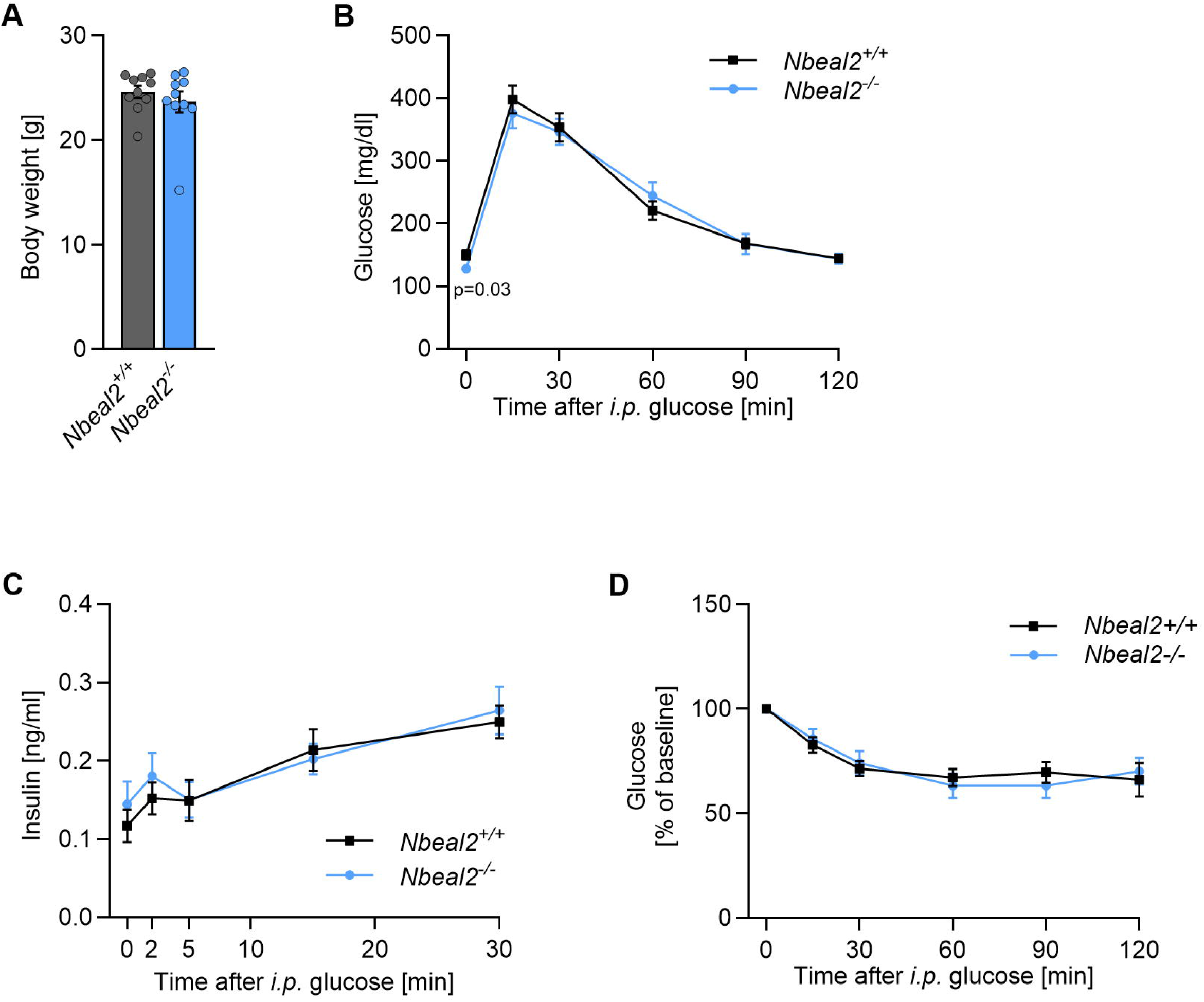
Nbeal2-deficiency does not affect glucose tolerance and glucose-stimulated insulin secretion. Body weight (**A**), glucose tolerance test (**B**), glucose-stimulated insulin secretion (**C**), and insulin tolerance test (**D**) of *Nbeal2*^-/-^ male mice fed a standard laboratory diet compared to respective littermate control animals. n=10 (A, B, D); n=6 (C). Each n represents the measurement of a sample from distinct mice. Mann-Whitney test. Data are presented as mean ± SEM.

### 3.2. Munc 13-4 deficiency results in a defect in glucose-stimulated insulin secretion

To address the effect of DGs on glucose homeostasis, we utilized *Unc13d*-deficient mice. *Unc13d* encodes Munc13-4, a protein required among others for DG exocytosis from platelets. In the absence of *Unc13d*, the secretion of DGs is almost completely blocked, and the release of AGs is markedly reduced [7]. In order to limit *<u>Unc13d</u>* <u>deficiency to hematopoietic stem cells</u>, we transplanted bone marrow from *Unc13d*-deficient mice into 6-week-old wild-type BL6 recipient mice (*Unc13d*^*BMC-/-*^). Control animals were generated by transplantation of bone marrow from wild-type littermates of *Unc13d*-deficient animals into BL6 recipient mice (*Unc13d*^*BMC*+/+^). After 4 weeks of recovery, we performed a glucose tolerance test. Notably, *Unc13d*^*BMC-/-*^ mice showed reduced glucose tolerance compared to control animals (Figure 2A). This was associated with reduced glucose-stimulated insulin secretion (Figure 2B). However, insulin sensitivity was not affected by the deletion of *Unc13d* (Figure 2C). Additionally, the body and organ weights of *Unc13d*^*BMC-/-*^ mice were not affected (Figures 2D and E). These data suggest that DG content mediates the influence of platelets on insulin secretion.

**Figure 2.**
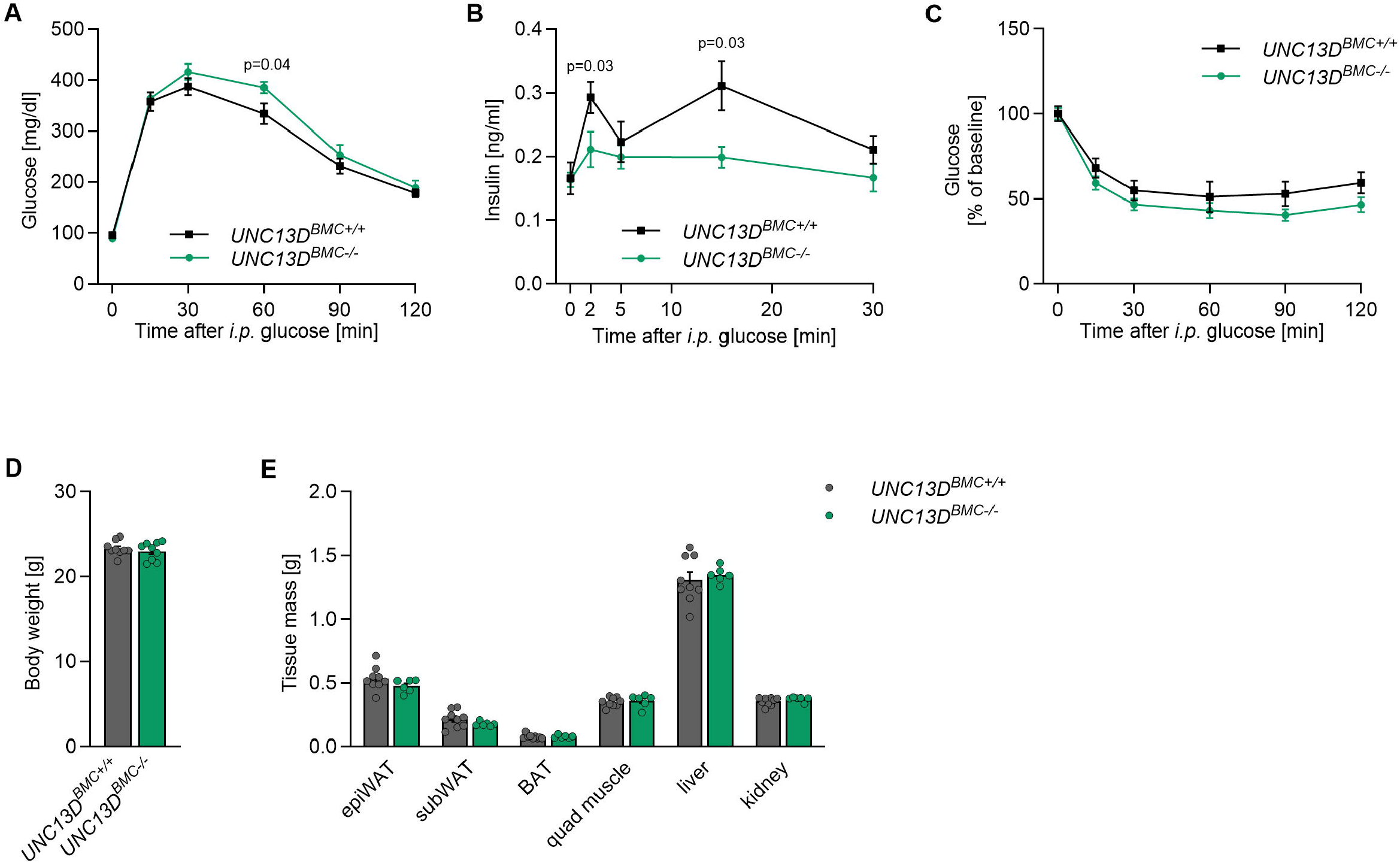
Unc13d-deficiency in bone marrow-derived cells results in glucose intolerance due to decreased glucose-stimulated insulin secretion. Glucose tolerance test (**A**), glucose-stimulated insulin secretion (**B**), insulin tolerance test (**C**), body weight (**D**), and tissue mass (**E**) of *UNC13D*^*BMC-/-*^ bone marrow chimeric male mice fed standard laboratory diet with respective littermate control animals. n=9 (A,C-D); n=8, *UNC13D*^*BMC+/+*^ and n=7, *UNC13D*^*BMC-/-*^ (B); n=9, *UNC13D*^*BMC+/+*^ and n=6, *UNC13D*^*BMC-/-*^ (E). Each n represents the measurement of a sample from distinct mice. Mann Whitney test. Data are presented as mean ± SEM.

### 3.3. Munc13-4 deficiency potentiates diet-induced glucose intolerance

Since mice with *Unc13d* deficiency in blood cells exhibit decreased glucose tolerance and insulin secretion, we utilized *Unc13d*-deficient mice in the bone marrow compartment (generated as described above) to explore the role of Munc13-4 in the development of diabetes. At 6 weeks of age (directly after bone marrow transplantation), mice were maintained on a high-fat diet (HFD). Importantly, *Unc13d*^*BMC-/-*^ mice showed reduced glucose tolerance after 20 weeks of HFD feeding (Figure 3A). Similar to mice on a normal diet, this was associated with impaired glucose-stimulated insulin secretion, while peripheral insulin tolerance was not affected by *Unc13d* deficiency in the bone marrow compartment (Figure 3B and C). These data indicate that Munc 13-4 regulates insulin secretion and glucose tolerance also in obese animals.

**Figure 3.**
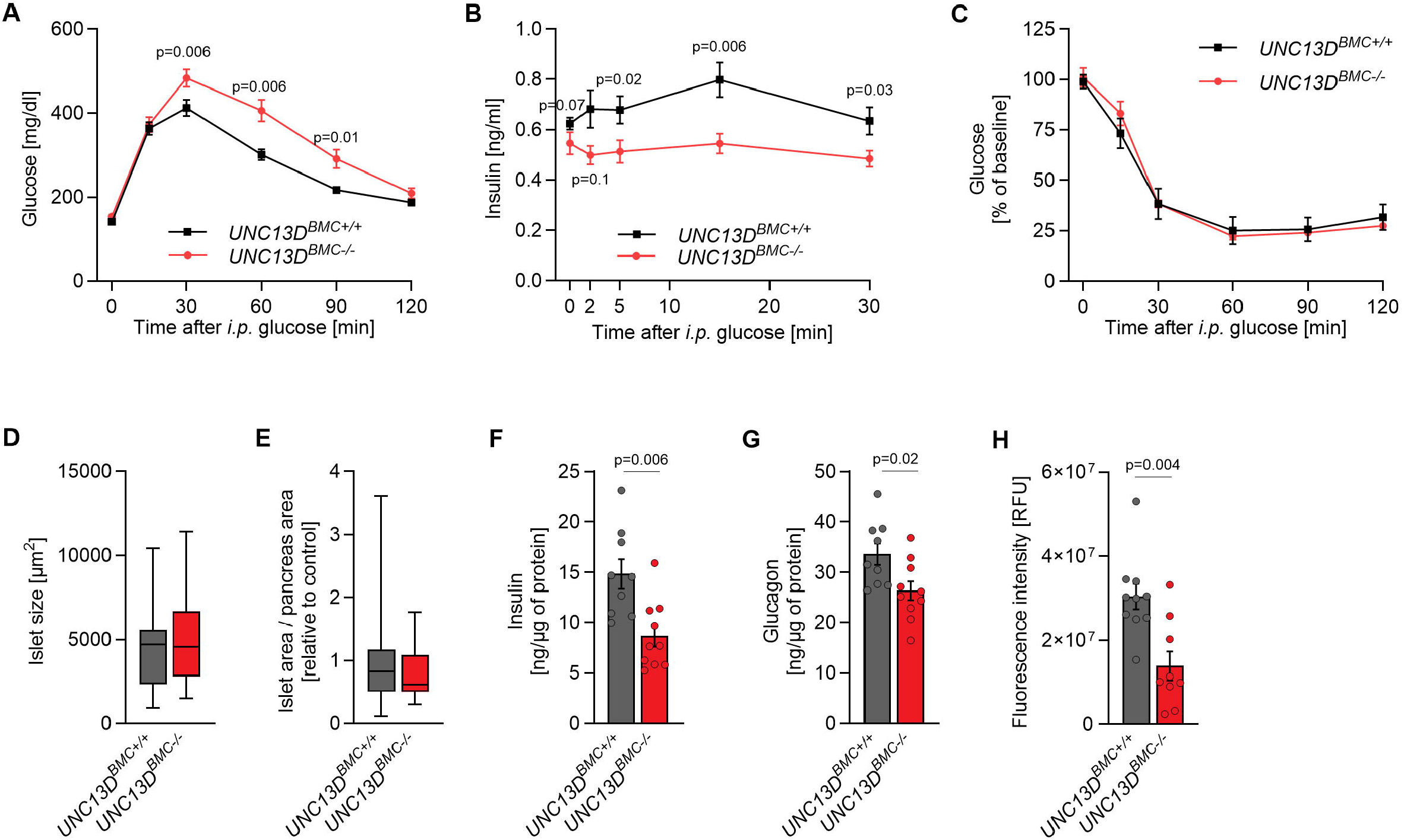
Unc13d-deficiency in bone marrow-derived cells results in impaired glucose homeostasis and decreased insulin and glucagon content in pancreas in obese animals. Glucose tolerance test (**A**), glucose-stimulated insulin secretion (**B**), insulin tolerance test (**C**), islet size (**D**), relative islet area (**E**), insulin (**F**) and glucagon (**G**) content in pancreas, and relative intensity of insulin staining in pancreatic islets (**H**) of *UNC13D*^*BMC-/-*^ bone marrow chimeric male mice fed high fat diet with respective littermate control animals. n=13 (A); n=10 (B); n=13, *UNC13D*^*BMC+/+*^ and n=14, *UNC13D*^*BMC-/-*^ (C); n=25, *UNC13D*^*BMC+/+*^ and n=31, *UNC13D*^*BMC-/-*^ (D); n=22, *UNC13D*^*BMC+/+*^ and n=27, *UNC13D*^*BMC-/-*^ (E); n=9, *UNC13D*^*+/+*^ and n=10, *UNC13D*^*BMC-/-*^ (F,G); n=10, *UNC13D*^*+/+*^ and n=9, *UNC13D*^*BMC-/-*^ (H). Each n represents the measurement of a sample from distinct mice (A-C, F-H) or an image of an islet (D) or pancreas (E). Mann Whitney test. Mann-Whitney test. Data are presented as mean ± SEM.

### 3.4. Munc 13-4 deficiency results in decrease in pancreatic insulin and glucose content without affecting β cell mass

Insulin secretion is tightly connected to pancreatic β cell mass. However, the pancreatic islet area and size were not affected by the deletion of *Unc13d* in blood cells (Figure 3D and E). Interestingly, total insulin content in the pancreas was markedly reduced in the absence of *Unc13d* (Figure 3F). Similarly, the content of glucagon, a hormone produced by pancreatic α cells that counteracts insulin action, was also reduced in the absence of *Unc13d* (Figure 3G). Moreover, the relative fluorescence intensity of insulin staining in pancreatic islets was significantly decreased in mice with *Unc13d*-deficiency (Figure 3H). These data suggest that Munc 13-4 protein in blood cells affects not only insulin release from pancreatic β cells but also mediate the number of insulin-positive cells as well as an overall production of pancreatic islet hormones in obese mice.

### 3.5. Munc 13-4 deficiency reduces fat deposition in obese HFD-fed mice

The weight of *Unc13d*^*BMC-/-*^ mice was monitored throughout HFD feeding. No significant changes in body weight were observed between *Unc13d*^*BMC-/-*^ mice and control animals. However, *Unc13d*^*BMC-/-*^ mice showed a trend towards reduced weight between the 18th and 24th weeks of HFD feeding (Figure 4A). Detailed analyses revealed a reduction in total fat mass and the weight of epigonadal as well as subcutaneous fat depots in *Unc13d*^*BMC-/-*^ mice (Figure 4B and C). Changes in fat deposition might be caused by altered food intake, energy expenditure, or malabsorption of digested food. Notably, food intake, respiratory exchange ratio, energy expenditure, and activity were not affected by *Unc13d* deficiency in blood cells (Figures 4D, E, F, and G). Altogether, these data indicate that Munc 13-4 protein in blood cells regulates fat deposition in obese mice without influencing food intake or energy expenditure.

**Figure 4.**
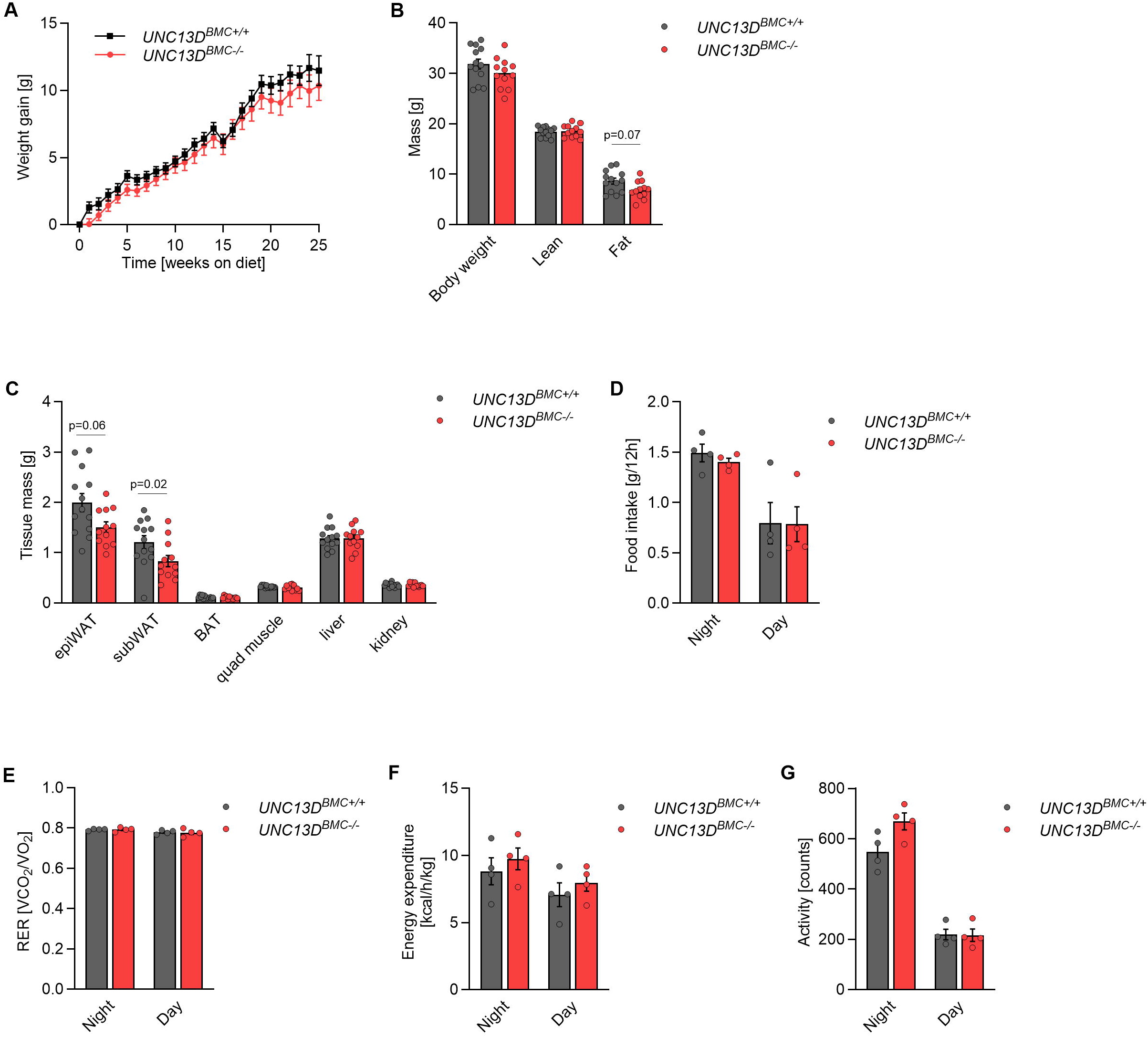
Unc13d-deficiency in bone marrow-derived cells reduces adiposity in mice fed a high-fat diet. Weight gain (**A**), body composition (**B**), tissue mass (**C**), food intake (**D**), respiratory exchange ratio (RER) (**E)**, energy expenditure (**F**), and activity (**G**) of *UNC13D*^*BMC-/-*^ bone marrow chimeric male mice fed a high fat diet compared to respective littermate control animals. n=13 (A); n=10 (B); n=13, *UNC13D*^*BMC+/+*^ and n=14, *UNC13D*^*BMC-/-*^ (C); n=14 (A); n=13, *UNC13D*^*BMC+/+*^ and n=12, *UNC13D*^*BMC-/-*^ (B,C); n=4 (D-G). Each n represents the measurement of a sample from distinct mice. Mann Whitney test. Data are presented as mean ± SEM.

## DISCUSSION

Our data indicate that the release of DGs by platelets is crucial for their action on pancreatic β cells, while α granules are dispensable for the induction of insulin secretion by these cells. A previous study identified a novel mechanism by which platelet-derived lipids, including 20-Hydroxyeicosatetraenoic acid (20-HETE), promote insulin secretion from pancreatic β cells [11]. Consequently, depletion of platelets, or the abrogation of platelet activation or adhesion, resulted in impaired glucose tolerance and glucose-stimulated insulin secretion [11]. Deletion of *Nbeal2* results in mice with mild thrombocytopenia [3-5]. However, our results indicate that the deletion of *Nbeal2* does not impair glucose tolerance or insulin secretion. One explanation for this discrepancy is that the previous study involved transient depletion of platelets, resulting in an almost complete lack of circulating platelets, whereas deletion of NBEAL2 results in only a moderate reduction in platelet count [3-5]. Moreover, a moderate decrease in platelet count might be potentially compensated by an increased production of substances promoting insulin secretion.

Conversely, defect in the release of DGs from platelets caused by the deletion of *Unc13d* [7], resulted in glucose intolerance and reduced glucose-stimulated insulin secretion. Importantly, *Unc13d*^*BMC-/-*^ mice fed a normal diet mimic the phenotype of animals with impaired platelet function [11]. A previous study indicates that platelets promote insulin secretion by releasing lipid-based substances. DGs contain multiple factors, including serotonin, ATP, ADP, and cations [2]. However, the presence of lipid-based substances in DGs has not been reported so far. This raises the question of whether 20-HETE and/or other active molecules required for insulin secretion are stored in DG and the ablation of DG release directly reduces the release of those substances or whether the blockage of DG secretion results in reduced platelet activation and consequently decreases the release of lipid-based substances (including 20-HETE).

In our research we utilized mice with *Nbeal2* global deletion and *Unc13d* deficiency in hematopoietic stem cells, thus described genetic modifications are not platelet specific. Despite the fact that observed phenomena could not be only due to changes in platelet degranulation, a comparison of our findings to previously described phenotypes of mice with defects in platelet function let us to conclude that the release of DGs, not AGs by platelets is crucial for their action on insulin secretion. Nevertheless, further investigation is required to answer the question of whether isolated platelets from *Nbeal2* and *Unc13d-*deficient mice themselves are able to potentiate insulin secretion from β cells.

Deletion of *Unc13d* also reduced glucose tolerance and insulin secretion in mice fed HFD to induce obesity. Previous studies indicated that mice deficient for Gαq and Gα13 specifically in platelets, which results in diminished activation of platelets by thrombin, ADP, or thromboxane A2 [15-17], presented unaltered glucose tolerance when fed HFD [11]. Similarly, pharmacological blockage of platelet activation with clopidogrel did not affect glucose tolerance in HFD-fed mice [11]. However, depletion of platelets in obese mice fed HFD resulted in reduced glucose tolerance [11]. Taken together, previous studies indicate that the effect of platelets on pancreatic β cells was largely abrogated by HFD feeding. These phenomena were initially explained by the fact that HFD feeding results in elevated levels of multiple lipid species [18], which saturate receptors for 20-HETE and other lipid derivatives at the level of pancreatic β cells. However, our current results challenge this concept. One possible explanation is that ineffective DG release might result in decreased insulin secretion by affecting mechanisms alternative to the previously proposed release of lipid-based substances. For instance, under HFD conditions, substances stored in DGs like serotonin, ATP, or ADP might influence β cell function independently of a saturation of the lipid-based signal. Interestingly, in HFD-fed animals deficient for *Unc13d*, we observed decreased insulin and glucagon content, which might suggest an alternative mechanism of β cell regulation by platelets under normal and HFD feeding conditions. Previous studies have emphasized the role of substances stored in DGs in platelets, particularly serotonin, in regulating pancreatic β cell function and other metabolic aspects of metabolism [19, 20]. However, the detailed mechanisms mediating platelet action on pancreatic β cells in HFD-fed mice need further clarification. Another possible explanation of the inconsistency of the phenotype of HFD-fed *Unc13d*^*BMC-/-*^ and previously described mice with impaired platelet function is that the observed phenomena is due to the fact that *Unc13d* deficiency is not platelet but blood cells-specific and defects in degranulation in other blood cells can contribute to the observed phenotype. This possibility is supported by data obtained from Jinx mice, that exhibit mutation in the Unc13D gene resulting in the absence of functional Munc13-4 protein and defective degranulation not only in platelets but also in immune cells [9, 21].

Interestingly, Okunishi *et al*. showed that HFD-fed Jinx mice are characterized by increased body weight and adiposity, as well as decreased glucose and insulin tolerance [22]. This is in contradiction to our data that shows no changes in body weight and insulin tolerance and decreased adipose tissue mass in Unc13D-deficient mice. The observed discrepancies may be due to the location of the mutation within a broader genomic region on chromosome 11, which encompasses multiple genes, and the phenotype of Jinx mice not necessarily is related only to a mutation in the Unc13D gene [21].

Our studies also showed that *Unc13d*-deficient mice fed HFD exhibit decreased adiposity. A previous study reported an increased abundance of platelets in adipose tissue from obese subjects. Moreover, this study indicates that platelets promote the differentiation of adipocytes [23]. Therefore, the release of DGs from platelets might promote the acquisition of new adipocytes and its defective degranulation could be responsible for lower adiposity. Additionally, as insulin promotes lipogenesis in adipose tissue, a decreased level of this hormone could contribute to the lower adiposity in *Unc13d*^*BMC-/-*^ mice [24]. Since our data indicate that food intake and energy expenditure are not affected by the deletion of *Unc13d*, proper degranulation in blood cells might be relevant for efficient nutrient absorption in the gastrointestinal tract. Alternatively, the composition of lean and fat mass could be changed without affecting overall energy absorption, as the total weight of animals is not affected by the deletion of *Unc13d*.

In conclusion, we demonstrated that the release of DGs, but not AGs by platelets promotes insulin levels and glucose homeostasis in mice.

## Abbreviations

20-HETE: hydroxyeicosatetraenoic acid
AGs: alpha granules
DGs: dense granules
HFD: high fat diet
Nbeal2: neurobeachin-like 2

## ACKNOWLEDGMENTS

This work was supported by the Dioscuri Centre of Scientific Excellence Grant number UMO-2018/01/H/NZ4/00002 – the program initiated by the Max Planck Society (Max-Planc-Gesellschaft), managed jointly with the National Science Centre in Poland (Narodowe Centrum Nauki) and mutually funded by the Polish Ministry of Science and Higher Education (Ministerstwo Nauki i Szkolnictwa Wyższego) and the German Federal Ministry of Education and Research (Bundesministerium für Bildung und Forschung) to G.S. and by the Deutsche Forschungsgemeinschaft (SFB1525, project number: 453989101 to D.S.).

## Notes

**Conflict of interest:** The authors have declared that no conflict of interest exists.

### Competing Interest Statement

The authors have declared no competing interest.

